# Arimoclomol as a potential therapy for neuronopathic Gaucher Disease

**DOI:** 10.1101/281824

**Authors:** Cathrine K. Fog-Tonnesen, Paola Zago, Erika Malini, Lukasz M. Solanko, Paolo Peruzzo, Claus Bornaes, Raffaella Magnoni, Nikolaj H. T. Petersen, Bruno Bembi, Andrea Dardis, Thomas Kirkegaard

## Abstract

Gaucher Disease (GD) is caused by mutations of the *GBA* gene which encodes the lysosomal enzyme acid beta-glucosidase (GCase). *GBA* mutations commonly affect GCase function by perturbing its protein homeostasis rather than its catalytic activity. Heat shock proteins (HSPs) are well known cytoprotective molecules with numerous functions in protein homeostasis and lysosomal function and their manipulation has been suggested as a potential therapeutic strategy for GD. The investigational drug arimoclomol, which is currently in phase II/III clinical trials, is a well-characterized HSP amplifier and has been extensively clinically tested. Importantly, arimoclomol efficiently crosses the blood-brain-barrier hereby presenting an opportunity to target the neurological manifestations of GD, which remains without a disease modifying therapy.

In the present study, we found that arimoclomol induced relevant HSPs such as ER-resident HSP70 (BiP) and enhanced the folding, maturation, activity and correct cellular localization of mutated GCase across a number of genotypes including the common L444P and N370S mutations in primary cells from GD patients. These effects where recapitulated in a human neuronal model of GD obtained by differentiation of multipotent adult stem cells. Taken together, these data demonstrate the potential of HSP-targeting therapies in GCase-deficiencies and strongly support the clinical development of arimoclomol as a potential first therapeutic option for the neuronopathic forms of GD.

**Summary:** These studies provide proof-of-concept for the development of the Heat shock protein amplifier, arimoclomol, as a potential therapy for neuronopathic Gaucher disease as arimoclomol enhances folding, maturation, activity and correct localization of GCase in neuronopathic and non-neuronopathic Gaucher disease models.

## Introduction

Gaucher disease (GD) is one of the most prevalent human metabolic storage disorders belonging to the group of lysosomal storage diseases (LSDs)(Grabowski 2008). It is primarily caused by autosomal recessive mutations in the *GBA* gene leading to deficiency of the lysosomal enzyme acid beta-glucosidase (GCase, EC 3.2.1.45). To date, more than 460 *GBA* mutations, the majority being missense mutations, have been identified (http://www.hgmd.cf.ac.uk)(Hruska et al. 2008). The mutations commonly lead to increased protein misfolding, premature degradation and abnormal chaperone recognition, which in turn lead to reduced GCase function. GCase dysfunction leads to the accumulation of its substrate glucosylceramide (GlcCer) and other sphingolipids, including glucosylsphingosine (GlcSph) causing cellular dysfunction and subsequent clinical manifestations primarily in the central nervous system (CNS), visceral and bone systems(Platt 2014).

GD is clinically divided in visceral type I (GD1), acute neuronopathic type 2 (GD2) and sub-acute neuronopathic type 3 (GD3) forms, although the age of onset and the phenotypic expression of the disease is variable(Grabowski et al. 2015). Visceral involvement in GD includes liver and spleen enlargement and dysfunction, as well as the displacement of normal bone marrow by storage cells causing anemia, thrombocytopenia and bone disease. Although GD1 is considered a non-neuropathic form, there is increasing evidence that neurological symptoms (i.e. Parkinson’s syndrome, tremors, peripheral neuropathy) become a prominent part of the pathology as the disease progresses(Bembi et al. 2003; Rosenbloom et al. 2011; Chetrit et al. 2013; Chérin et al. 2010; Devigili et al. 2017). GD2 is very rare (1% of cases) and is associated with a severe and rapid neurodegeneration, leading to an early death, usually before the second year of life(Weiss et al. 2015). In GD3, visceral and biochemical signs are similar to GD1 and the first symptoms are usually due to peripheral involvement. The involvement of the CNS generally appears later and includes oculomotor apraxia, ataxia, epilepsy and mental deterioration(Tylki-Szymańska et al. 2010). Currently, two types of treatments are available for GD1: enzyme replacement therapy (ERT) and substrate reduction therapy (SRT) but there are no approved disease-modifying treatments for the neurological forms of the disease.

The molecular mechanisms involved in the neurodegenerative process in GD are not fully elucidated but the disease pathology ultimately stem from the loss of function of GCase. Mutations in the gene are numerous with the majority being missense mutations that affect the correct folding and maturation of the protein, but do not completely abrogate the catalytic activity(Hruska et al. 2008). In GD, misfolded GCase is retained in the endoplasmic reticulum (ER), from where it is retro-translocated back to the cytosol to be eliminated by the ubiquitin proteasome pathway(Bendikov-Bar et al. 2011). This process, known as ER-associated degradation (ERAD), impede mutant GCase from reaching the lysosome, leading to compromised lysosomal GCase activity and function. In addition, the presence of misfolded GCase triggers the unfolded protein response (UPR) and interferes with the degradation of other proteins(Maor et al. 2013). The loss of GCase function leads to accumulation of metabolites such as GlcCer and GlcSph, expansion of a compromised, destabilized lysosomal compartment, inflammation and loss of neurons potentially involving the Rip3k pathway(Platt et al. 2016; Platt 2014; Vitner et al. 2014).

Members of the HSP70 family of molecular chaperones include HSP70 encoded by *HSPA1A* and BiP/GRP78/HSP70-5 encoded by *HSPA5* (Daugaard et al. 2007), which have been shown to be important for lysosomal and GCase function(Wang et al. 2011; Kirkegaard et al. 2016; Kirkegaard et al. 2010; Ingemann & Kirkegaard 2014; Jian et al. 2016). Studies of GCase, and its most prevalent missense mutations for non-neuronopatic (N370S) and neuronopathic (L444P) GD, have demonstrated that it binds to HSP70 and HSP90 which in concert with cochaperones such as TCP1 guide the enzyme through to either correct folding and lysosomal activity, or to the ubiquitin proteasome pathway for degradation(Lu et al. 2011; Yang et al. 2014; Yang et al. 2015; Jian et al. 2016).

Several lines of evidence indicate that it is possible to at least partially rescue GCase folding and function by strategies aimed at increasing the levels of cellular molecular chaperones such as HSP70 or through the use of small molecules acting as chemical chaperones(Khanna et al. 2010; Sun et al. 2011; Maegawa et al. 2009; Mu et al. 2008; Yang et al. 2013; Yang et al. 2014; Yang et al. 2015; Lu et al. 2011).

Arimoclomol, a small molecule amplifier of HSP70 and other HSP chaperones, has emerged as a potential therapeutic agent for several LSDs including Niemann-Pick Disease Type C (NPC)(Kirkegaard et al. 2016) and is currently being evaluated in a phase II/III trial for NPC (ClinicalTrials.gov identifier NCT02612129). Importantly, previous clinical studies of other neurodegenerative diseases, such as amyotrophic lateral sclerosis (ALS) and sporadic inclusion body myositis (sIBM), have demonstrated a safety profile for arimoclomol compliant with chronic use, as well as penetrance to the CNS and signs of efficacy(Cudkowicz et al. 2008; Lanka et al. 2009; Ahmed et al. 2016). Based on the clinical safety-profile, the CNS-penetrable ability and the HSP-inducing mechanism of action, arimoclomol may present a first-in-class treatment paradigm for GD patients – particularly patients with currently untreated neurological symptoms. We therefore investigated the effect of arimoclomol on the stability, localization and enzymatic activity of GCase across a broad range of genotypes in primary cultured GD fibroblasts and in a human neuronal model of GD obtained through differentiation of multipotent adult stem cells.

## Results

### Arimoclomol increases the quantity and ER to golgi maturation of mutated GCase in primary GD patient fibroblasts

We obtained a panel of primary skin-derived fibroblast cell lines from individuals diagnosed with GD covering the major genotypes and from three healthy donors. Sequencing of the *GBA* gene confirmed the presence of the reported pathogenic mutations of *GBA* in all GD cell lines (Table 1). We also identified the presence of a T369M variant in the widely used control fibroblast cell line GM05659 [WT/T369M]. T369M does not cause GD in homozygous carriers, but may be associated with an increased risk of developing Parkinson’s disease(Alcalay et al. 2015; Mallett et al. 2016).

**Table 1.**
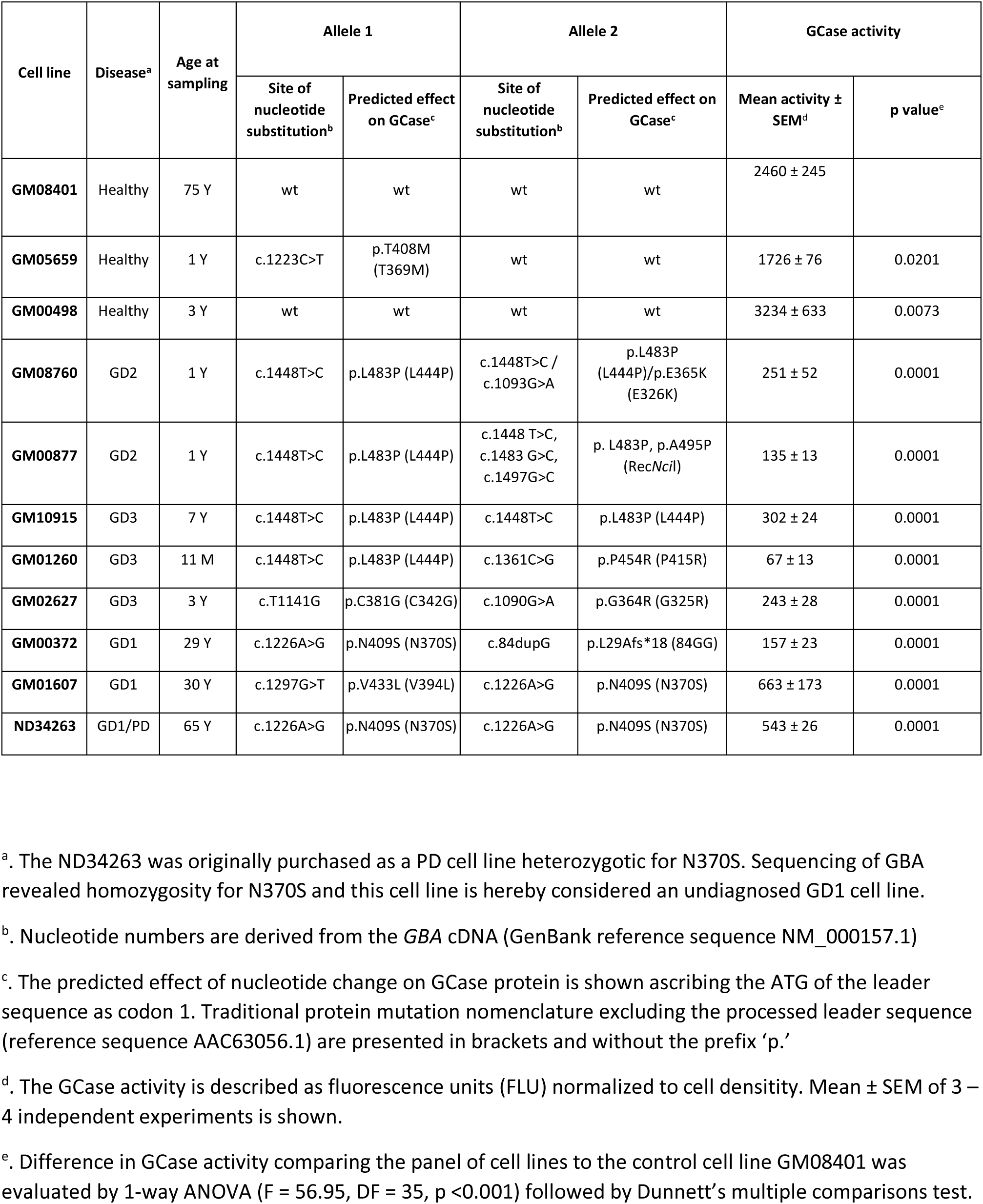
Characteristics of primary human fibroblast cells lines.

Comparison of mRNA, protein and GCase activity levels across the WT and GD primary patient fibroblasts demonstrated no correlation between the level of *GBA* mRNA and disease (Figure 1a), whereas neuronopathic GD (nGD) cell lines showed a profound decrease in the GCase protein level compared to WT fibroblasts (Figure 1b). The GD1 cell line GM00372 [N370S/1-bp ins 84G] also displayed a clear decrease in the amount of GCase protein, which is likely caused by the premature stop codon being produced by the 1-bp ins 84G allele (Beutler et al. 1991), whereas two other non-neuronopathic GD cell lines GM01607 [N370S/V394L] and ND34263 [N370S/N370S] had only slightly reduced GCase protein levels (Figure 1b).

**Figure 1.**
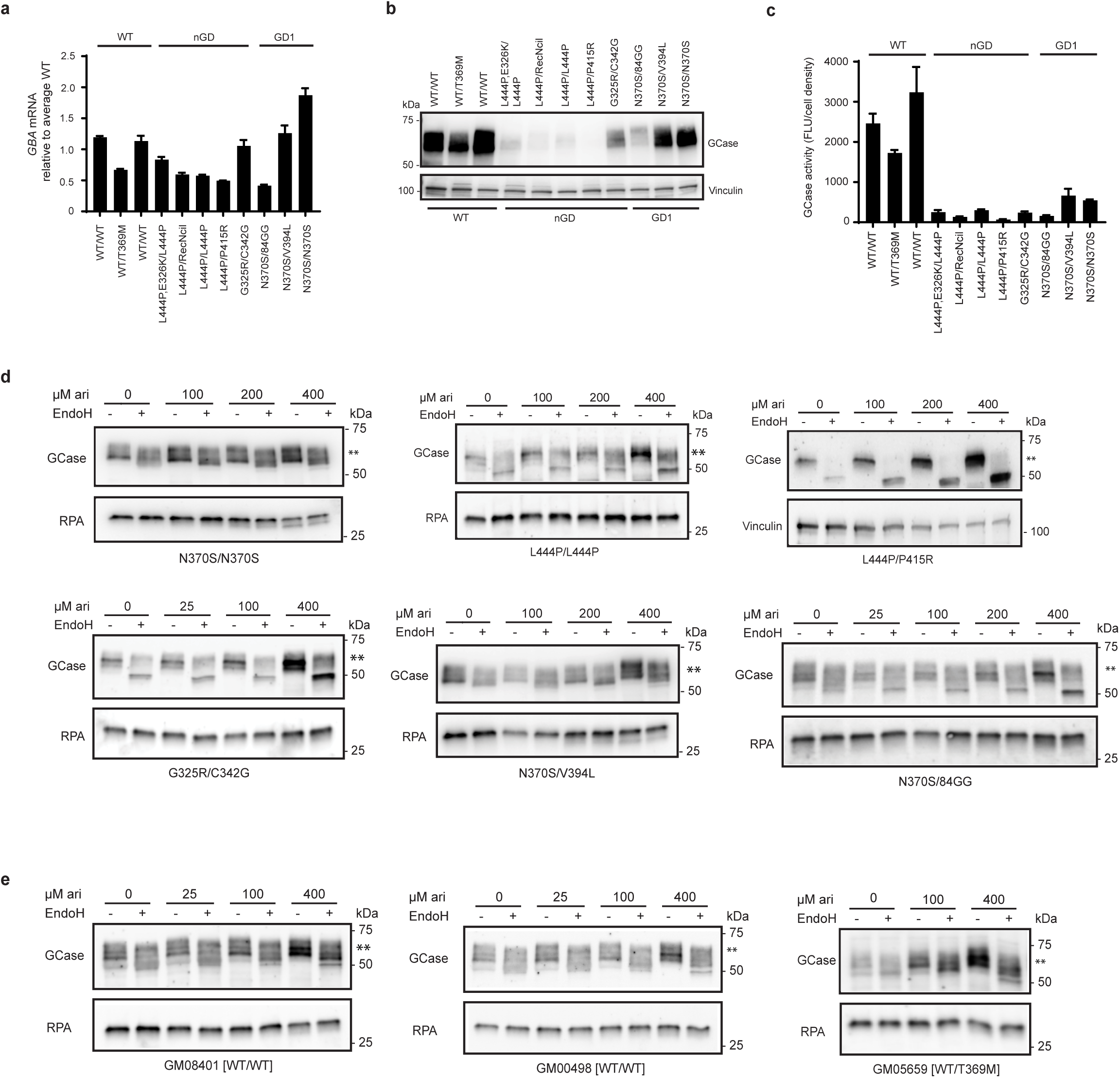
Arimoclomol increases the quantity and ER to golgi maturation of mutated GCase in primary GD patient fibroblasts. a) Relative level of *GBA* mRNA expression in primary GD fibroblasts normalized to the average expression levels in WT cells. b) WB analysis of GCase protein levels in primary GD fibroblasts and WT control cell lines. Vinculin was used as loading control. c) Basal level of GCase activity (Fluorescence units (FLU) normalized to cell density) in the primary GD fibroblasts and WT control cell lines. Data shown as mean + SEM of 3-4 experiments/cell line. d-e) WB analysis of GCase in (d) primary GD fibroblasts or (e) WT control cell lines treated with the indicated concentrations of arimoclomol for 5 days. Lysates were subjected to EndoH-digestion analysis and the EndoH-resistant fraction is marked by **. RPA or Vinculin served as loading control for these experiments. WBs are representative of 3 independent experiments.

Analysis of basal GCase activity showed reduced activity of the mutated GCase in all the GD cell lines investigated (Figure 1c).

Arimoclomol is a well-described co-inducer of the Heat shock response, which includes the amplification of HSP70 chaperones (Vígh et al. 1997; Kalmar et al. 2012). We first tested if arimoclomol amplifies and prolongs the expression of HSP70 and BiP in primary GD cells. Time and dose evaluations as well as qPCR and western blots confirmed that arimoclomol significantly increases HSP70 and BiP levels in neuronopathic and non-neuronopathic GD primary fibroblasts (Supplementary Figure 1a-c).

We then investigated the folding and ER to golgi transition of mutant GCase variants with or without arimoclomol treatment. Cell lysates were digested with Endoglycosidase H (EndoH), an endoglycosidase that specifically cleaves high mannose (>4 mannose residues) but not mature N-glycan complexes, allowing differentiation between immature glycoproteins that have not reached the mid-Golgi (EndoH-sensitive) and mature glycoproteins (EndoH-resistant)(Maley et al. 1989).

Arimoclomol dose-dependently increased the amount of GCase in primary GD fibroblasts across all tested genotypes, including neuronopathic and nonneuronopathic associated mutations (Figure 1d). Treatment with arimoclomol also increased EndoH resistant levels of GCase for all genotypes including the most common non-neuronopathic and neuronopathic mutations (N370S and L444P, respectively).

EndoH digestion analysis of WT GCase demonstrated that WT GCase is subject to some degree of EndoH digestion as reported previously (Sawkar & Kelly 2006; Bendikov-Bar et al. 2011) and that arimoclomol could also augment WT GCase levels (Figure 1e).

#### Arimoclomol increases residual GCase activity in GD cells

Having demonstrated a beneficial effect of arimoclomol on the maturation of mutant GCase protein in primary GD fibroblasts we next evaluated the effect of arimoclomol on GCase activity. The GM10915 [L444P/L444P] cell line was treated with 50 – 800 µM arimoclomol for 1 – 5 days and GCase activity was measured using 4-MUG as substrate. A time- and dose-dependent increase in GCase activity was seen in cells treated with arimoclomol corresponding to the observed amplification of HSP70 family members (Figure 2a and Supplementary figure 1a-c). Longer exposure (28 days) at lower concentrations of arimoclomol resulted in similar increases of GCase activity (Figure 2b).

**Figure 2.**
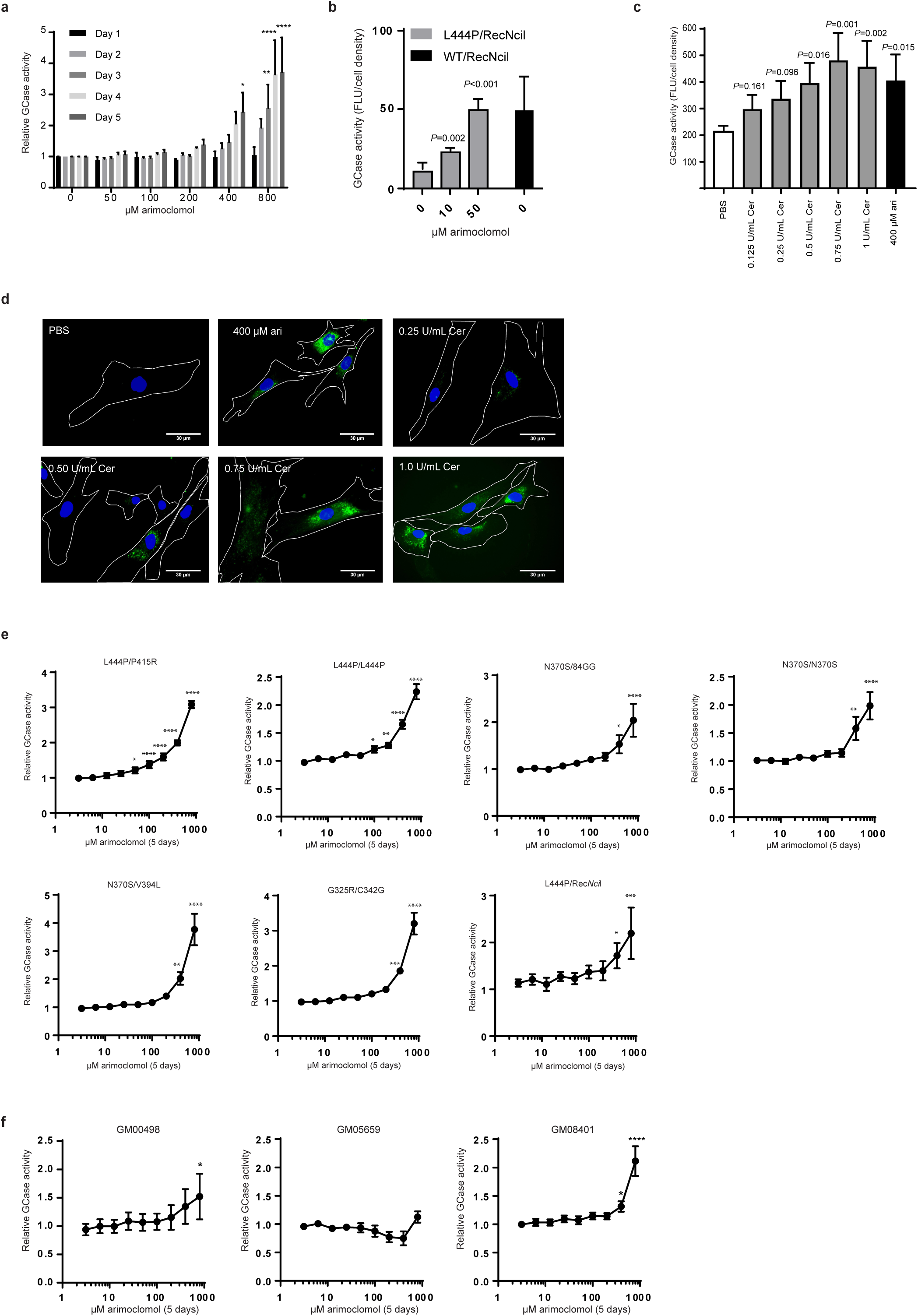
Arimoclomol augments residual GCase activity in primary GD patient fibroblasts. a) GCase activity in GM10915 cells treated with 50 – 800 µM arimoclomol for 1 – 5 days. Relative GCase activity is shown as fold change compared to vehicle-treated cells for each time point. Data are mean + SEM of 3 independent experiments. The effect of each arimoclomol concentration was evaluated against vehicle within the same day using a 2-way RM ANOVA model with the interaction arimoclomol concentration*day as fixed effect. Multiplicity was adjusted using Dunnett’s method. (*<0.05, **<0.01, ***<0.001, ****<0.0001). b) GCase activity in GM00877 (L444P/RecNcil) and GM00878 (WT/RecNcil) cells treated with arimoclomol for 4 weeks. Data are shown as mean + SD, n=3-6. c) Effect of 400 µM arimoclomol (ari) and 0.125 – 1 U/mL Cerezyme (Cer) on GCase activity levels in GM10915 cells treated for 5 days. Data are shown as mean + SEM, n=4. Effects of treatments were compared using 1-way RM ANOVA with multiplicity adjusting using Dunnett’s method. d) Representative images of GCase labeled with ABP-green in GM10915 (L444P/L444P) cells treated with vehicle (PBS), 400 µM arimoclomol or 0.25 – 1.0 U/mL cerezyme for 5 days. e) Relative GCase activity in primary GD patient fibroblasts treated with arimoclomol for 5 days. f) GCase activity in WT fibroblast cell lines treated with the indicated concentrations of arimoclomol for 5 days. For e-f Data are shown as mean ± SEM of 3-4 independent experiments/cell line. Statistical analysis was done using a 1-way RM ANOVA model. Multiplicity was adjusted using Dunnett’s method (*<0.05, **<0.01, ***<0.001, ****<0.0001).

To qualify the increases in GCase activity observed with arimoclomol we examined the levels of GCase activity obtainable with the current standard of care therapy for non-neuronopathic GD, enzyme replacement therapy, in the form of recombinant beta-glucocerebosidase (Cerezyme®). We treated L444P/L444P cells with Cerezyme^®^ or arimoclomol for 5 days. The cellular uptake and activity of Cerezyme^®^ from the cell culture medium was confirmed using activity-based probes (ABPs)(Witte et al. 2010) (Figure 2d). A dose-dependent increase in GCase activity was obtained for Cerezyme®, with maximum obtainable activity levelling out at 0.5-1.0 U/mL in both systems. The increase in GCase activity obtained by 400 µM arimoclomol was comparable to the maximum attainable activity levels with Cerezyme^®^ (Figure 2c-d).

We then proceeded to investigate the effect of 5-days treatment with arimoclomol on residual GCase activity across our panel of primary GD fibroblasts. In line with the observed effects on HSP70 induction and GCase maturation, arimoclomol significantly increased residual GCase activity across all genotypes including both neuronopathic and nonneuronopathic alleles (Figure 2e). We also observed significant increases in two of three WT cell lines (Figure 2f).

#### Arimoclomol improves the correct lysosomal localization of GCase in GD cells

To asses if the rescued GCase reaches the correct intracellular localization and to validate the observed improvement in maturation and activity increases, we took advantage of the highly specific fluorescent ABPs which allow labeling of active GCase molecules in cells(Witte et al. 2010). WT, GM01607 [N370S/V394L], GM02627 [G325R/C342G] and GM10915 [L444P/L444P] cells were treated with 400 µM arimoclomol for 5 days. Imaging of fixed cells showed clear punctuate high intensity GCase localization in WT cells, whereas GD cell lines displayed decreased labeling (Figure 3a). The GCase labelling intensity was increased in arimoclomol-treated primary GD patient fibroblasts, with a clear lysosomal distribution pattern. The primary image analysis was corroborated by automated image quantification demonstrating a significant fluorescence intensity shift towards the WT profile in the arimoclomol treated GD fibroblasts compared to untreated controls (Figure 3b-d).

**Figure 3.**
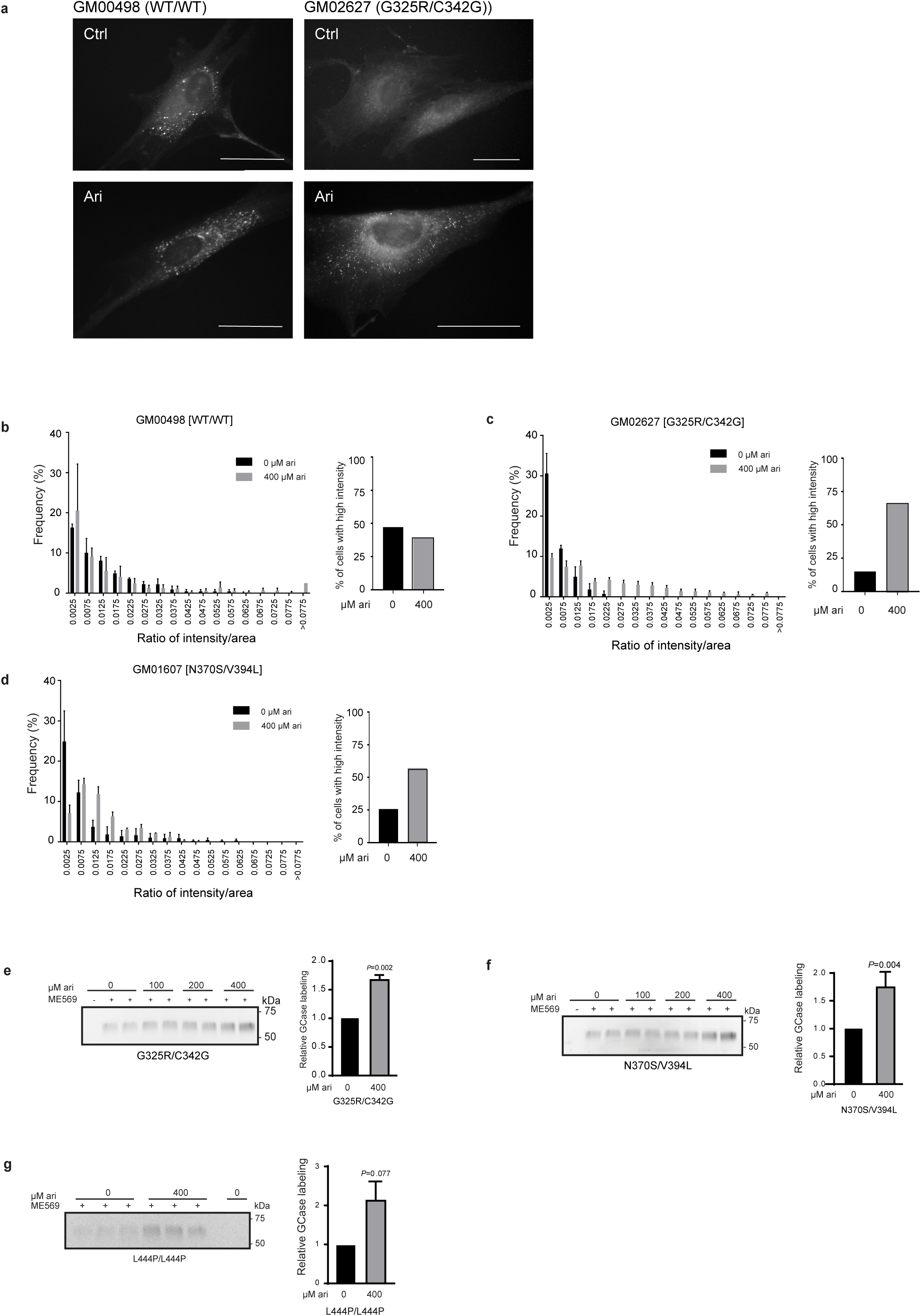
Arimoclomol increases lysosomal localization and activity of GCase in GD fibroblasts. a) Representative images of primary WT or GD patient fibroblasts treated with arimoclomol for 5 days and labeled with green fluorescent ABPs (left). Scale bars = 30µm. b-d) Image analysis quantification of active GCase labeling. The quantification of labeling is shown as the frequency distribution of the ABP labeling intensity per area in grouped intervals, n=3, >100 cells analysed per replicate. e-g) Gel quantification of ABP-labelling of active GCase. Representative fluorescent gel images of cell lysates labeled with ABP-cy5 ME569. Cells were treated with arimoclomol for 5 days and each concentration was evaluated in duplicate or triplicate samples. The quantification of ABP-labeling is shown in the right panel as mean + SEM, n=3-4. The effect of arimoclomol was analyzed by RM oneway-ANOVA. Multiplicity was adjusted using Dunnett’s method.

We further quantified the effect of arimoclomol treatment on active GCase labeling in GD cells by using SDS-page resolved cell lysates incubated with ABP (Figure 3e-g). The amount of active, labeled GCase was significantly increased by arimoclomol for both non-neuronopathic and neuronopathic genotypes.

#### Arimoclomol increases GCase activity in neuronal-like cells from GD patients

We next evaluated the effect of arimoclomol in a human neuronal model of GD, obtained by induced neuronal differentiation of multipotent adult stem cells (MASCs) isolated from neuronopathic GD patients(Bergamin et al. 2013a). MASC-derived neuronal cultures were established from a healthy donor and from neuronopathic GD patients carrying two different genotypes: F213I/L444P and L444P/L444P. In addition, MASCs were obtained from one non-neuronopathic GD patient (Table 2). *GBA* sequencing identified this GD individual as a compound heterozygous for the common N370S mutation and a variant not previously reported: c.516C>A that results in a codon change from tyrosine 133 (TAC) to a stop codon (TAA). However, the analysis of the *GBA* mRNA isolated from the patient’s cells showed that only the allele carrying the normal cytosine in position 516 was expressed (Supplementary Figure 2a). These results suggest that the Y133* mutation leads to the expression of an unstable transcript resulting in either no or very little truncated GCase protein. We characterized the surface immunophenotype of the MASCs (Supplementary Table 2) as previously described and observed no major differences between MASCs from GD patients and healthy donors(Bergamin et al. 2013b). The differentiation of early passage MASCs (Passage 1-3) to neuronal-like cells was ascertained by immunostaining of the neuronal markers NeuN and tubulin beta 3(TUBB3) (Figure 4a, b). The presence of arimoclomol during differentiation did not impact the neuronal differentiation (Figure 4a, b).

**Table 2.** Characteristics of MASCs.

| Patient | Disease | Age at onset | Age at sampling | Allele 1 |  | Allele 2 |  | GCase activity neuronal like cells |  |
| --- | --- | --- | --- | --- | --- | --- | --- | --- | --- |
| | | | | Site of nucleotide substitution<sup>a</sup> | Predicted effect on GCase<sup>b</sup> | Site of nucleotide substitution<sup>a</sup> | Predicted effect on GCase<sup>b</sup> | Mean activity ± SEM<sup>c</sup> | p value<sup>d</sup> |
| 1 | GD1 | 23 | 33 | c.1226A>G | p.N409S (N370S) | c.516C>A | p.Y172* (Y133*) | 7.95 ± 0.32 | <0.0001 |
| 2 | GD3 | 3 | 30 | C754T>A | p.F252I (F213I) | c.1448T>C | p.L483P (L444P) | 8.20 ± 0.77 | <0.0001 |
| 3 | GD3 | 3 | 58 | c.1448T>C | p.L483P (L444P) | c.1448T>C | p.L483P (L444P) | 6.16 ± 1.31 | <0.0001 |
| 4 | GD3 | 6 | 14 | c.115+1G>A (IVS2+1G>A) | p.(E10_S38del G39Vfs51*) | c.680A>G | p.N227S (N188S) | 20.95 ± 2.68 | <0.0001 |
| 5 | healthy donor | | 32 | WT | WT | WT | WT | 316.13 ± 10.89 | |

<sup>a</sup>. Nucleotide numbers are derived from the *GBA* cDNA (GenBank reference sequence NM\_000157.3)
<sup>b</sup>. The predicted effect of nucleotide change on GCase protein is shown ascribing the ATG of the leader sequence as codon 1. Traditional protein mutation nomenclature excluding the processed leader sequence (reference sequence AAC63056.1) are presented in brackets and without the prefix 'p.'
<sup>c</sup>. The GCase activity (nmol/mg/h) is reported as mean ± SEM, n=2-4
<sup>d</sup>. Differences in GCase activity in neurons from GD individuals and a healthy donor were evaluated by 1-way ANOVA ((F(4,11) = 1289, p <0.0001)) followed by Dunnett's multiple comparisons test. P-values are reported as multiplicity adjusted p-values

**Figure 4.**
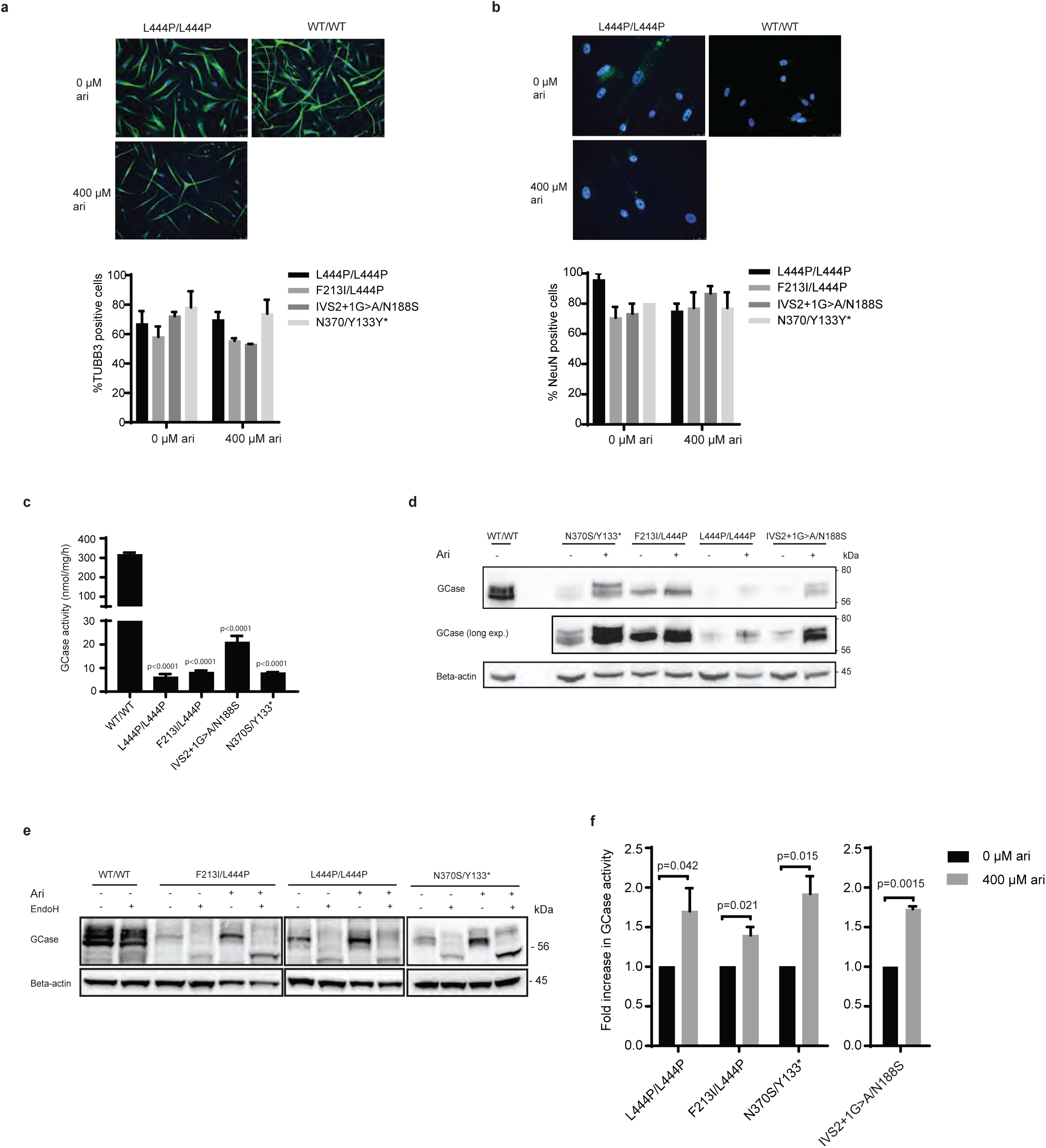
Arimoclomol increases GCase expression, lysosomal localization and activity of GCase in human neuronal-like GD cells. a - b) Representative images of differentiated human neuronal like cells stained for expression of a) TUBB3 and b) NeuN. The quantification of % positive cells derived from four GD patients with the indicated genotypes are shown below. c) GCase activity in human neuronal-like cells derived from healthy donors (WT/WT) or GD individuals with the indicated *GBA* mutations. Data are shown as mean + SEM, n=2-4/cell line. The difference in GCase activity between WT and GD neuronal-like cells were analyzed by oneway-ANOVA and multiplicity was adjusted by Dunnet’s method. d) Representative WB analysis of GCase levels in human neuronal-like cells derived from healthy donors (WT/WT) or GD individuals treated with vehicle or arimoclomol (400 µM). Beta-actin was used as loading control. e) EndoH-digestion assay of neuronal-like cells treated vehicle or 400 µM arimoclomol. A representative WB from one out of 2 experiments is shown. f) GCase activity in in GD-derived human neuronal cells treated with vehicle or 400 µM arimoclomol for 9 (L444P/L444P, F213I/L444P, N370S/F133*) or 11 (IVS2+1G/N188S) days. Data are shown as fold increase in arimoclomol-treated cells compared to vehicle-treated cells and represent the mean + SEM, n=4-6. Statistical significance was assessed by unequal variance t-test.

The level of GCase activity was analyzed in the final stage of neuronal differentiation. The GD-derived neurons displayed severely diminished activity of GCase compared to healthy donor derived cells (Figure 4c, Table 2). In line with the results obtained in GD patient fibroblasts, low levels of GCase activity were associated with reduced levels of GCase protein in neuronally differentiated GD cells compared to healthy controls (Supplementary Figure 2b).

We proceeded to investigate the effect of arimoclomol on the level, maturation and activity of GCase in GD-patient derived neuronal cell cultures (Figure 4d-f). Treatment with arimoclomol augmented both the level and EndoH resistant fraction of GCase in both neuronopathic and non-neuronopathic GD patient derived neuronal cells (Figure 4d, e). In line with the WB and EndoH-assay results, treatment with arimoclomol also resulted in a corresponding significant increase in GCase activity in neuronal cells from both neuronopathic and non-neuronopathic GD patients (Figure 4f).

## Discussion

Our results demonstrate that arimoclomol is a heat shock protein amplifying small molecule that may be useful for the treatment of Gaucher disease including its neuronopathic forms which have no approved treatments available.

Arimoclomol amplifies the production of disease mechanism-relevant molecular chaperones of the HSP70 family and improves mutant GCase maturation and function across major neuronopathic and non-neuronopathic genotypes in both human primary GD fibroblasts as well as in a neuronal model of the disease.

As arimoclomol is a clinically enabled compound already in phase II/III clinical trials for Niemann-Pick disease type C and sporadic inclusion body myositis (Clinicaltrials.gov identifiers NCT02612129 and NCT02753530, respectively), the data reported herein provide preclinical proof-of-concept for the investigation of arimoclomol’s therapeutic value in Gaucher disease.

Arimoclomol amplifies the production of HSP70 family members which have been implicated in the proper folding and chaperoning of GCase as well as in maintenance of lysosomal integrity during cellular stress(Kirkegaard et al. 2010; Kirkegaard et al. 2016; Lu et al. 2011; Yang et al. 2014; Yang et al. 2013; Yang et al. 2015; Ingemann & Kirkegaard 2014; Mu et al. 2008). Studies of the induction of HSPs by small molecules have until now made use of agents that provide proof-of-principle but due to their cytotoxic nature are not suited for development for chronic diseases(Mu et al. 2008; Ingemann & Kirkegaard 2014; Kirkegaard 2013). These studies have however, revealed several HSP-dependent mechanisms regulating the processing of GCase. Studies in cellular models of GD have shown that HSP-induction by agents such as celastrol and MG-132 result in elevated GCase activity through increases in both cytosolic and ER-resident chaperones of the HSP70 family, through a process dependent on the UPR responsive transcription factors Ire1, ATF6 and PERK(Mu et al. 2008). Other studies have corroborated these findings in GD cells, by showing that celastrol upregulates HSP70 and its associated cochaperone BCL2-associated athanogene 3 (BAG3), interrupting the HSP90-dependent degradation of GCase(Yang et al. 2014; Yang et al. 2015). The role of HSP70 in the regulation of GCase activity has been further expanded by recent studies suggesting that HSP70 is recruited directly to GCase by progranulin (PGRN) acting as a critical cochaperone(Jian et al. 2016). In addition, several other HSP-inducing agents such as HDAC-, proteasome- and HSP90 inhibitors as well as L-type Ca-channel blockers have demonstrated that manipulation of the HSP-systems can improve the folding and activity of mutant GCase(Yang et al. 2013; Lu et al. 2011; Ingemann & Kirkegaard 2014; Mu et al. 2008; Wang & Segatori 2013). Interestingly, parts of the mechanism of how HDAC inhibitors elicit their GCase rescuing effect might be ascribed to their role as HSP90 inhibitors, as the HDAC inhibitors vorinostat and LB-205 both bind to the middle domain of HSP90, resulting in less recognition of misfolded GCase and simultaneous upregulation of the HSP70 chaperones involved in refolding(Yang et al. 2013). Studies of vorinostat in NPC has further expanded the knowledge on how HDAC inhibitors rescue misfolded lysosomal proteins through modulation of HSPs by a mechanism likely involving inhibition of HDAC1(Pipalia et al. 2011). This is supported by data demonstrating that Heat Shock Factor 1 (HSF1), the major transcription factor for HSP70 and other HSPs, specifically interacts with HDAC1 and HDAC2 to regulate gene expression during heat shock(Pipalia et al. 2011; Fritah et al. 2009).

Arimoclomol holds particular promise for neuronopathic GD as it is a well-tolerated, CNS-penetrant molecule currently in a Phase II/III clinical trial for another neurodegenerative LSD, NPC(Cudkowicz et al. 2008; Lanka et al. 2009) (Clinialtrials.gov identifier NCT02612129). Through its mechanism of increasing the expression of multiple HSPs, arimoclomol could potentially target GD and its CNS related events at other levels besides the HSP system’s capacity to improve the folding, maturation and activity of GCase as reported herein. By amplifying cellular levels of HSP70, other HSP70 lysosome-specific cytoprotective activities such as improving the function of other sphingolipid degrading enzymes and protection against lysosomal destabilization may be achieved, as has been recently demonstrated for a number of related sphingolipid storage diseases(Kirkegaard et al. 2010; Kirkegaard et al. 2016; Zhu et al. 2014; Nakasone et al. 2014; Ingemann & Kirkegaard 2014). The reported capacity of HSP70 to protect against lysosomal membrane permeabilization and lysosomal cell death pathways may be particularly interesting aspects of HSP70 amplification in GD as the storage metabolite glucosylsphingosine has been shown to initiate lysosomal dysfunction and cell death(Folts et al. 2016). Interestingly, it has recently been demonstrated that neuronal cell death in a model of GD(Vitner et al. 2014) is necroptotic and involves the receptor-interacting protein Kinase-3 (RIPK3) pathway which is regulated by the HSP70 cochaperone CHIP (Carboxyl terminus of HSP70-interacting protein)(Seo et al. 2016). Further, one of the hallmarks of necroptosis is lysosomal membrane permeabilization, a cellular event which HSP70 has been demonstrated to protect against in several disease models including sphingolipid storage diseases(Platt 2014; Kirkegaard et al. 2010; Kirkegaard et al. 2016; Nylandsted et al. 2004; Zhu et al. 2014; Bivik et al. 2007; Jaattela et al. 1992; Gyrd-Hansen et al. 2006; Hwang et al. 2005; Kirkegaard & Jäättelä 2009).

Although much remains to be understood about the molecular mechanisms leading to GD and in particular its neurological manifestations, it is clear that the cytoprotective properties of the Heat shock proteins, in particular HSP70 and its cochaperones, converge with the pathogenesis of GD at several critical levels.

Based on these observations and the data herein, we suggest that arimoclomol constitute a potential disease-modifying first-in-class compound for the treatment of Gaucher disease, in particular neuronopathic GD which is currently without efficacious treatment options.

## Methods

### Cell culture and drug treatment

All human fibroblast cell lines used for this study were obtained from Coriell Biorepositories and cultured under standard cell culture conditions (37 °C and 5 % CO_2_) in DMEM supplemented with non-essential amino acids (NEAA), 1% Pen-Strep and 12% FCS. They were passaged 1-2 times/week with a split ratio of 1:2 or 1:3. Cells were used for experiments around passage 16-26 where no signs of replicative senescence were observed (visual inspection).

Cells were treated with vehicle (PBS) or arimoclomol-citrate (BRX-345) dissolved in PBS for the indicated time and medium containing fresh compound was added every 2-3 days. For experiments with Cerezyme®, cells were treated for 5 days with medium replenishment every 2-3 days. Fresh medium containing compounds was also added on the day before visualization by ABPs.

### *GBA* sequencing

DNA was purified using the DNeasy Blood and Tissue kit (# 69504, Qiagen) according to manufacturer’s instructions. For each sample, the whole genomic *GBA* sequence (GenBank J03060.1) was PCR-amplified in two overlapped fragments of 2870 bp and 4492 bp, using primers designed to selectively amplify the gene and not the homologous pseudogene as previously described (Koprivica et al. 2000). Amplification products were purified using the QIAquick Gel Extraction Kit (Qiagen), quantified by Quant-iT PicoGreen dsDNA assay (Life technologies) and then processed using a Nextera XT DNA sample preparation kit (Illumina) to generate pair end libraries and sequenced on an Illumina MiSeq using an Illumina MiSeq Reagent kit v2 (MS-102-2002) 300 cycles as described (Zampieri et al. 2017).

All SNV and indel information are output in variant call format (VCF). Annotation of SNV was performed with wANNOVAR (http://wannovar.usc.edu/).

Sanger sequencing was performed to confirm exonic variants identified by massive sequencing and to exclude the presence of the recombinant allele Rec delta 55 in all samples as described in (Malini et al. 2014).

Mutations are described as recommended, considering nucleotide +1 the A of the first ATG translation initiation codon (http://www.hgvs.org/mutnomen/). Nucleotide numbers are derived from the *GBA* cDNA (GenBank reference sequence NM_000157.1). Traditional protein mutation nomenclature does not start with the first ATG codon due to a processed leader sequence (reference sequence AAC63056.1). These mutations are presented without ‘p.’ in the mutation name.

Sanger sequencing of *GBA* mRNA from cells carrying the N370/Y133* genotype: Total RNA was extracted from cell lines using Rneasy mini Kit (Qiagen). First-strand cDNAs were synthesized by Advantage RT-for-PCR Kit (Clontech, Mountain View, CA, USA) and random hexamer primers. Reverse transcriptase-PCR (RT-PCR) was performed using the following specific *GBA* primers: 5’acctgcatccttgtttttgtttagtggatc 3’ and 5’ gactctgtccctttaatgcccaggctgagc 3’. Then a fragment of 450 bp was obtained by nested PCR and analyzed by automated sequencing (ABI Prism 3500xl genetic analyzer, Applied Biosystems, Foster City, CA, USA).

### Quantitative real-time PCR

Total RNA was extracted using the RNeasy mini Plus kit (Qiagen) according to manufacturer’s instructions. cDNA was prepared using the QuantiTect RT kit (Qiagen). The primers used for quantitative real-time PCR (qPCR) are listed in Supplementary Table 1. qPCR was performed by using PowerUp SYBR green (Applied Biosystems) on the QuantStudio 3 system (Thermo Scientific) with the following settings: 50 °C 2 min, 95 °C 2 min. Then 40 cycles of 1 s at 95 °C, 30 s at 60 °C. Analysis was performed with the QuantStudio Design and Analysis software v. 1.4 (Thermo Scientific). Relative gene expression was determined using the δδCt method normalizing to the average of the house keeping genes *TUBB, HPRT* and *RPLPO*. Relative gene expression was calculated as fold change relative to control sample.

### Western blotting and EndoH assay

Cells were collected and lysed in 25 mM KPi buffer, pH 6.5 + 0.1% Triton X-100 including a cocktail of protease inhibitors (Roche). Protein concentration was determined using with the bicinchoninic acid protein assay (BCA) kit (Pierce). 10-20 µg total protein/sample was used for WB of protein levels or used for EndoH-treatment according to manufacturer’s instructions (New England Biolabs). Samples boiled in Laemmli sample buffer were subjected to SDS-PAGE using the TGX gel system (Bio-Rad). After transfer to a nitrocellulose membrane (Trans-Blot Turbo, Bio-Rad), the membranes were stained briefly with Ponceau S (Sigma-Aldrich), and subsequently blocked in 5 % skim-milk in PBS + 0.1 % tween (PBS-T). Incubation with primary antibodies (1:500 to 1:2000 dilution) was performed on parafilm-coated glass plates overnight at 4 °C. After washing in PBS-T membranes were incubated 1 h with secondary antibody (Vector Laboratories) diluted 1:10,000 in 5 % skim milk in PBS-T. The blots were developed using SuperSignal™ West Dura Extended Duration Substrate (Life technologies) and visualized using a G-box system (Syngene).

Antibodies used were: anti-GBA (# WH0002629M1, Sigma-Aldrich), anti-Vinculin (# V9131, Sigma-Aldrich), anti-GRP78 (BiP) (# sc-13968, Santa Cruz Biotechnology), anti-RPA (# sc-28709, Santa Cruz Biotechnology), anti-Beta actin (# A2066, Sigma-Aldrich).

### GCase activity

For fibroblasts, cells were seeded in 96 well plates and treated in biological triplicate with various concentrations of arimoclomol for the indicated time. Medium was replenished every 2-3 days. GCase activity was measured using 4-Methylumbelliferyl β-D-glucopyranoside (4-MUG) (Sigma-Aldrich) as substrate at pH 4.0 with overnight incubation at 37 °C and 5 % CO_2_. The reaction was stopped with glycine, pH 10.8 and the released 4-methylumbelliferone (4-MU) fluorophore was measured (excitation 365 nm, emission 445 nm) with VarioScan Flash (Thermo Scientific). For normalization to cell number, a parallel plate was stained with 0.1% crystal violet, dissolved in 1 % SDS and absorbance (570 nm) was measured. GCase activity was calculated as Fluorescence Units (FLU) normalized to cell density. For each experiment, the fold change (FC) of compound-treated versus mock-treated cells was calculated and the data reported as mean FC ± SEM of at least 3 independent experiments. Initially, by addition of 1 mM Conduritol B Epoxide (CBE) (Sigma-Aldrich), non-lysosomal GBA2 activity in fibroblasts was found to be less than 3 % of the total fluorescence signal.

GCase activity in neuronal-like cells was measured in cell lysates. Cells were lysed in H_2_O and sonicated for 15 s. Protein concentration was determined with the Bradford reagent (Bio-Rad, Hercules, CA, USA). 7 µg protein was assayed for GCase activity by incubation with 7.5 mM 4-MUG for 3 h at 37 °C. Stop buffer pH 10.7 was added and the released 4-MU fluorophore was quantified using a Gemini XPS Spectrometer (Molecular Devices, Sunnyvale, CA, USA) (excitation length: 340 nm; emission: 495 nm).

### ABP labelling and quantification

Fibroblasts were collected and lysed in 25 mM KPi buffer, pH 6.5 + 0.1% Triton X-100 including a cocktail of protease inhibitors (Roche). Protein concentration was determined using with the BCA kit (Pierce). For labelling, equal amount of protein (usually 20 µg) was incubated with cy5-labelled ABP ME569 in 150 mM McIlvaine buffer, pH 5.2 for 1 h at 37 °C. Samples were denatured with 4× Laemmli buffer, boiled for 5 min at 98 °C and resolved by SDS-PAGE using the TGX gel system (Bio-Rad). Fluorescence was detected by G:BOX ChemiXR5 (Syngene) with red LED lightning modules/705M filter. The amount of labelled GCase was quantified using GeneTools v.4.03.01.0 (Syngene).

For visualization, fixed cells were incubated with ABP MDW933 (green) and images were acquired using the AXIO (Zeiss) equipped with AxioCam MRm (LED ex 470nm, em 500-550nm). Images were analysed using freeware program Fiji (an image processing package to ImageJ, https://fiji.sc/). Every acquired image was corrected for shading and autofluorescence and segmented to visualize a single cell per field of view (FOV). For each of FOVs, dynamic threshold was set to encapsulate cell’s boarders and calculate mean intensity. Collection of results was processed using GraphPad prism ver. 7, where entries were scanned for outliers and analysed for frequency distribution as a function of intensity/area per bin

### MASC generation and differentiation

Human skin-derived multipotent adult stem cells (MASCs) were obtained from skin biopsies from healthy donors and patients affected by GD, who were under observation at the Regional Centre for Rare Diseases. Written consent was obtained from the subjects or from caregivers or guardians on behalf of the minors involved in the study.

MASC purification and differentiation protocols were previously described (Bergamin et al. 2013a). Briefly, after 3 passages in selective medium, MASCs were detached and stained with the following primary conjugated antibodies: CD13, CD49a, CD49b, CD49d, CD90, CD73, CD44, CD45, human leukocyte antigen-D related (HLA-DR), CD34, and CD271 (BD Biosciences, Franklin Lakes, NJ, USA); CD105 and kinase insert domain receptor (KDR; Serotec, Oxford, United Kingdom); and CD133 (Miltenyi Biotec, Bergisch Gladbach, Germany). The percentage of cells expressing all the antigens was determined by fluorescence-activated cell sorting (FACS) analysis (CyAn; Beckman Coulter, Brea, CA, USA). Properly conjugated isotype-matched antibodies were used as negative controls.

For neurogenic differentiation, MASCs were plated in medium containing DMEM-HG with 10 % FBS (STEMCELL Technologies) (N1 medium). After 24 h the DMEM-HG was replaced with fresh medium supplemented with 1 % of B27 (Invitrogen), 10 ng/ml human EGF (Peprotech) and human 20 ng/ml bFGF (Peprotech) (N2 medium) for 5 days. Thereafter, cells were incubated for 72 h in DMEM-HG supplemented with 5 μg/ml insulin, 200 μM of indomethacin and 0.5 mM IBMX (all from Sigma-Aldrich) without FBS (N3 medium). The actual differentiation was determined by analyzing the expression of the neuron specific markers, NeuN and tubulin b3 (TUBB3).

For immunofluorescence assays, cells were fixed for 15 min in 4 % (w/v) paraformaldehyde in PBS and then permeabilized in 0.3 % Triton X-100 in PBS for 5 min on ice. After blocking with 2 % BSA in PBS, cells were incubated overnight at 4 °C with the primary antibodies raised against NeuN (Millipore) or tubulin beta 3 (Covance, Inc., Princeton, NJ, USA). Cells were then washed and incubated with Alexa Fluor 555 or 488 labelled secondary antibody for 1 h at 37 °C. Cell nuclei were stained by Vectashield Mountain Medium with DAPI (Vector Laboratories, Inc., Burlingame, CA, USA). Images were obtained with a live cell imaging dedicated system consisting of a Leica DMI 6000B microscope connected to a Leica DFC350FX camera (Leica Microsystems, Wetzlar, Hessen, Germany).

For treatment, vehicle or 400 μM arimoclomol was included in the N2 medium, replenished after 3 days and added to the N3 medium for a total treatment time of 9 days.

### Statistical analysis

GraphPad Prism v.7.03 was used for statistical analysis. Unless otherwise stated, repeated measures (RM) ANOVAs were calculated using a mixed model for repeated measures with treatments/timepoints as fixed effects and experiment as repeated random effect. Multiplicity was adjusted using Sidak-Holm or Dunnett’s method. Welch-Satterthwaite t-tests were used to correct for unequal variance.

## Author contributions

T.K. conceived and supervised the study. C.F.T., A.D., C.B., N.H.T.P. and T.K. designed the experiments with the contribution of P.Z, E.M, L.M.S, R.M. and B.B.

C.F.T, L.M.S, C.B, M.M.E and A.K. performed the experiments with human fibroblasts. P.Z., E.M. and P.P. performed the experiments with MASCs. E.M. and A.D. performed sequencing analysis. C.F.T., N.H.T.P., A.D. and T.K. wrote the manuscript.

## Acknowledgements

We thank Johannes M. Aerts and Hermen S. Overkleeft for ABPs and valuable scientific discussions. We also thank Mingshu Meng Eriksen, Anja Koustrup and Barbara Toffoletto for their technical assistance with regard to fibroblast experiments (MME & AK) and assistance in the immunophenotype characterization of MASCs (BT).

TKJ, CFT, NHTP, CB, LMS and RM are employees of Orphazyme A/S. TKJ is a Founder of and hold shares in Orphazyme A/S. Orphazyme A/S funded this work. The authors have no additional financial interests.

## Supplementary material

The supplementary material consists of 2 supplementary figures and 2 supplementary tables.

1. **Supplementary Figure 1. Arimoclomol increases and prolongs the HSR in GD fibroblasts.**
2. **Supplementary Figure 2. Sequencing analysis of genomic DNA and mRNA of MASCs carrying the N370/Y133* genotype**
3. **Supplementary Table 1:**
4. **Supplementary Table 2:**

**Supplementary Figure 1. Arimoclomol increases and prolongs the HSR in GD fibroblasts.**

a) Expression of *HSPA1* and *HSPA5* in GM10915 cells without heat shock (0 h timepoint) or at the indicted recovery times after 1 h at 41.5 °C. Prior to heat shock, cells were treated with vehicle or 400 µM arimoclomol for 72 hrs. Each time point was measured in 2 – 4 independent experiments and data are shown as mean + SEM. The effect of arimoclomol was evaluated against control at each time point by a two-way ANOVA and multiplicity was adjusted by Holm-Sidak’s method (*<0.05, **<0.01, ***<0.001, ****<0.0001). b) Expression of *HSPA1A* (HSP70) or *HSPA5* (BiP) in GM10915 cells treated with vehicle (0 µM arimoclomol) or 400 µM arimoclomol for the indicated time. Expression levels, relative to the 0 hour timepoint, are shown as mean ± SEM of 3 independent experiments. Statistical analysis was done by 2-way RM ANOVA and multiplicity was adjusted using Sidak-Holm. c) WB analysis of BiP in GM02627 and GM01607 cells treated with the indicated concentrations of arimoclomol for 5 days. Vinculin was used as loading control. WBs are representative of 3 independent experiments.

**Supplementary Figure 2. Sequencing analysis of genomic DNA and mRNA of MASCs carrying the N370/Y133* genotype**

a) Analysis of genomic DNA showed the presence of the c.516C>A mutation in heterozygosis (left panel), while sequencing of the cDNA extracted from patient’s cells showed the absence of the mutation at position 516 of the cDNA indicating that expression of the mutated allele cannot be detected. b) WB analysis of basal GCase protein levels in the final neuronal differentiation state of WT and GD-derived MASCs.

